# Establishment and maintenance of two sand fly colonies (*Phlebotomus perniciosus* and *Phlebotomus papatasi*) to implement experimental infections

**DOI:** 10.1101/2023.07.19.549665

**Authors:** Jorian Prudhomme, Lison Laroche, Anh Nguyen Hoàng Quân, Justine Fournier, Idris Mhaidi, Aloïs Berhard, Mireille Killick-Kendrick, Anne-Laure Bañuls

**Affiliations:** UMR MIVEGEC (Université de Montpellier - IRD – CNRS), Institute of Research for Development, Montpellier, France; INTHERES, Université de Toulouse, INRAE, ENVT, Toulouse, France; University of Science and Technology of Hanoi (USTH), Hanoi, Viet Nam; 2 Place du Temple, 30440 Sumène, France

**Keywords:** Sand flies, Phlebotomus, insect rearing, life cycle, experimental infection

## Abstract

Sand flies are vectors of *Leishmania* and *Phlebovirus*, pathogens responsible for public health concerns in numerous countries. Sand fly colonies are essential for research on vector-pathogen interactions. Methods for establishing and maintaining sand fly colonies have been well described in the literature. However, very few colonies of these considerable vectors currently exist.

In order to encourage the creation of new colonies, we provide a timeline for researchers who want to implement colonies of *Phlebotomus perniciosus* and *Phlebotomus papatasi*. In this study, we set up colonies of both species to test and optimize their breeding conditions.

For *Ph. perniciosus*, we reached a stable production of 800 individuals per week, one year after the start of the colony. For *Ph. papatasi*, a stable production of 400 individuals per week was reached 11 months after establishing the colony. Moreover, we have successfully conducted experiments within a year of establishing the colonies.

In conclusion, our data on these two species showed that colonies could be established without difficulty with 1000-1500 eggs. It is possible to reach a stable weekly production of 800 individuals one year after starting the colony. We also describe colony settings and breeding recommendations to facilitate this procedure for other laboratories.

## Introduction

Sand flies are small hematophagous dipteran insects. There are more than 900 species of sand flies, and among these 98 are potential vectors of diseases (Maroli et al., 2013). They are vectors of *Leishmania* and *Phlebovirus*, and both these pathogens remain a public health problem in many countries (World Health Organization, 2018).

*Leishmania* can cause different clinical forms in humans, from asymptomatic or benign cutaneous forms, to visceral forms which are lethal without treatment. About 20 *Leishmania* species are pathogenic to humans (Ready, 2010). For example, *Leishmania infantum* is endemic around the Mediterranean basin and can cause cutaneous or visceral forms. The number of leishmaniasis cases and their distribution areas are still on the rise and many reports show that this disease is far from being eradicated (Antoniou et al., 2013; World Health Organization, 2018). Moreover, there is a re-emergence in old foci and emergence in new foci (Aoun & Bouratbine, 2014; Ozbilgin et al., 2019).

As for the genus *Phlebovirus*, it comprises about 70 viruses, including the Toscana virus, responsible for human meningoencephalitis in the Mediterranean region. This virus is the main cause of aseptic meningitis during the warm season in Spain, Italy and Southern France (Bichaud et al., 2014). Both pathogens, *Leishmania* and *Phlebovirus*, co-circulate in the Mediterranean basin (Moriconi et al., 2017).

Many physical and behavioral characteristics of these insects are still unknown (*e*.*g*., inter and intraspecific vectorial capacity, poorly known food preferences). Sand fly colonies for experimental infection are essential to conduct research on these vector insects and the pathogens they transmit. However, there are only few sand fly colonies in the world. In 2016, only around 100 colonies, with 33 different species, were found worldwide (Lawyer Phillip et al., 2016).

The objective of our study was to establish *Phlebotomus perniciosus* and *Ph. papatasi* colonies. *Phlebotomus perniciosus* is the main vector of *Le. infantum* in Southwestern Europe (Bruschi & Gradoni, 2018). *Phlebotomus papatasi* is one of the major vectors of *Leishmania major* in the Old World (Ready, 2013). Methods to establish and maintain sand fly colonies have been well described in the literature (Volf & Volfova, 2011; Lawyer P et al., 2017; Molina et al., 2017). However, there are few colonies of these two species although they are easy to breed and important vectors.

In order to encourage the creation of new colonies of these sand flies, we provide a timeline for researchers who want to implement sand fly colonies of the species *Ph. perniciosus* and *Ph. papatasi*. In this study, in 2020, we established, in IRD Vectopôle (Montpellier, France), colonies of both species to test and optimize their breeding conditions.

## Materials and methods

### Ethical statement

Rabbit blood draws performed in the context of this study were approved by the Animal Care and Use Committee named “Comité d’Ethique pour l’Expérimentation Animale Languedoc Roussillon n°36” under protocol number 2018022712203932. Rabbits, coming from the animal facility at IRD, were not subjected to anesthesia, analgesia or sacrifice. The chicks come from the Experimental Infectiology Platform of Nouzilly (INRAE Centre Val de Loire, France). They were killed according to the Directive 2010/63/EU (Appendix IV) appropriate to the species. The chicks were dead before the use of tissue samples for experimental infection, in accordance with article R214-89 of the French “Code rural et de la pêche maritime”, Section 6.

### Starting colonies

We received sand fly eggs in November and March 2020 for *Ph. perniciosus* and *Ph. papatasi*, respectively. To start the sand fly colonies, we used eggs from well-adapted laboratory colonies of *Ph. perniciosus* (Laboratorio de Entomología Médica, Instituto de Salud Carlos III, Madrid, Spain) and *Ph. papatasi* (Department of Parasitology, Charles University, Prague, Czech Republic). The eggs were shipped in rearing pots by our collaborators Pr. Ricardo Molina and Pr. Petr Volf. Our colonies were started, in IRD Vectopôle (Montpellier, France), with approximatively 1,500 and 1,000 eggs from *Ph. papatasi* and *Ph. perniciosus*, respectively.

#### Recommendations

To start a colony, we recommend to use eggs when the species are available from other laboratories for the following reasons: 1) transportation is easier, 2) this method does not require blood feeding and, 3) it makes it possible to collect data from the first generation. However, it is also possible to use individuals collected from the field (Volf & Volfova, 2011).

### Sand fly rearing

Sand flies were maintained under standard conditions (26 ± 1°C, 80% relative humidity, 14h/10h light-dark cycle). For colony maintenance, we used protocol described by Volf & Volfova (2011), Lawyer P et al. (2017) and Molina et al. (2017), with some modifications described below.

Adults were maintained in large fabric-net holding cage (30×30×30cm) and fed with 50% glucose solution (protocol in Supp. Table S1). Blood feeding was done with glass feeders, filled with rabbit blood and covered with chicken skin membranes. Five-days-old chicks were used as they present a thin skin which allows efficient blood feeding (protocol in Supp. Table S2). The blood in the feeders was maintained at a 37°C constant temperature by a water bath with external circulation. Feeding was performed for five-six hours with a regular manual blood mix.

After 24 hours, blood-fed females were transferred in clear plastic containers filled with 1cm layer of Paris plaster. These pots were stored in plastic boxes containing humidified sand (Volf & Volfova, 2011). After hatching, larvae were fed with a matured mixture of 40% rabbit feces, 40% rabbit chow, and 20% mice chow (protocol in Supp. Table S3).

We performed blood feeding once a week for *Ph. papatasi*. For *Ph. perniciosus*, three blood feedings per week were done for the first generation (colony amplification), once a week for the second generation (due to COVID-19 lockdown measures), twice a week from the third generation to seventh generation (colony amplification), and once a week from the seventh generation onwards (colony routine maintenance).

#### Recommendations

Using glass feeders allows longer blood feeding and consequently increases blood-fed female numbers. To facilitate feeding, stimulate females every 30 minutes by breathing on the cage. Moreover, if the purpose of the colony is to perform experimental infections, the same protocol can be used in BSL2 or BSL3 laboratories, with the exception of stimulation by breathing (Vaselek et al., 2020). During experimental infection, adding males (10%) will improve the engorgement rate (Volf & Volfova, 2011).

We recommend three blood feedings a week for the two first generations (colony amplification) and then once a week (routine maintenance).

### Monitoring of sand fly colonies

We monitored the breeding data daily: mortality rate, mean blood feeding rate, development time, average generation time and productivity of the colony. Ranges of minimum developmental times were obtained by observing the first appearance of each stage. The recording of these parameters was done every day for the first generation, twice a week for the second generation (due to COVID-19 sanitary restrictions) and then four times a week (routine maintenance). The goal was to achieve the minimum of care without impacting the colony.

We used these parameters to determine the duration of each development stage duration, colony productivity and acclimatization.

#### Recommendations

In the early stages of the colony, it is imperative to come every day to check the survival rate, as well as to identify and solve possible problems as soon as they appear. A daily check of the larvae also prevents an overly rapid fungi proliferation which can be lethal for the larvae.

For standardized data collection and tracking, we recommend the use of a detailed file available in Supp. Table S4. The follow-up of these parameters makes it possible to plan experiments and to check for any extended generation time (*i*.*e*., the first sign of an ailing colony) (Volf & Volfova, 2011).

### Use of sand fly colony

Colony productivity determination makes it possible to define when and how many individuals are available for experiments. In our case, we performed insecticide resistance tests (World Health Organization, 2016) on *Ph. papatasi* and experimental infections (Vaselek et al., 2020) on *Ph. perniciosus*. Indeed, the setting up of sand fly colonies has the following objectives: 1) to ensure the continuation of the colony and 2) to conduct suitable experiments (experimental infections or insecticide resistance testing).

#### Recommendations

The colony size has to be monitored and it is essential to determine the number of individuals that can be sacrificed for the experiments. Indeed, to maintain the colony, no more than a third of its size should be taken (Lawyer P et al., 2017). Females under three to four days old do not feed and females aged more than ten days have a lower survival rate after blood feeding (Volf & Volfova, 2011). We used females aged from four to eight days (females produced over the week). As such, it is important to know the weekly production but also how much time is necessary for production to meet the needs of the laboratory.

### Statistical analysis

Statistical analyses of developmental times in different generations were performed using a nonparametric Kruskal-Wallis test followed by a post-hoc test (*i*.*e*., Mann-Whitney tests with Bonferroni correction), using R software version 3.1.0 (R Development Core Team, 2018).

## Results and discussion

### Colony establishment

The colony is currently in its eighth generation for *Ph. papatasi*, and in its tenth generation for *Ph. perniciosus*. The number of individuals in the colonies according to time is detailed in Figure 1.

**Figure 1.**
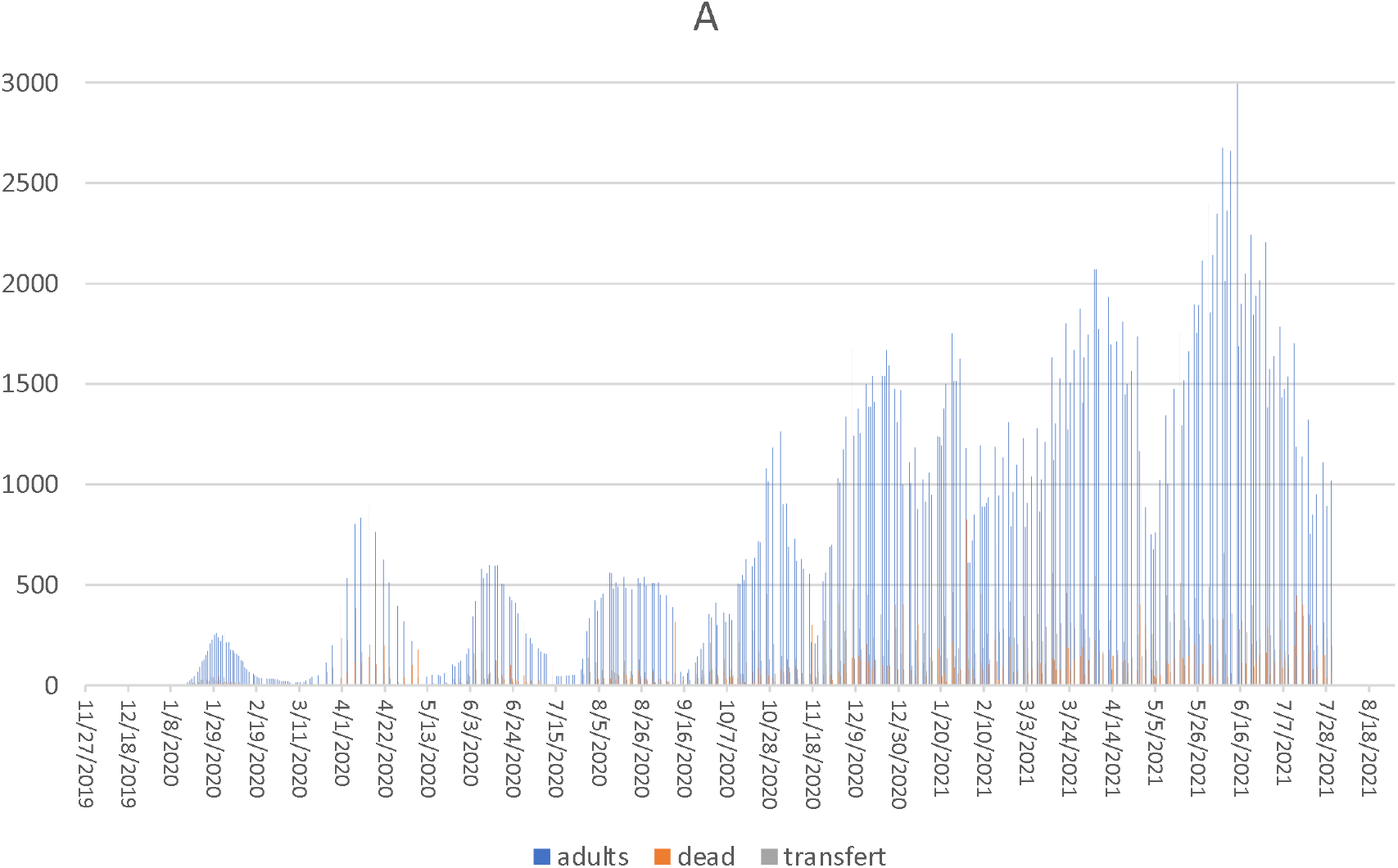
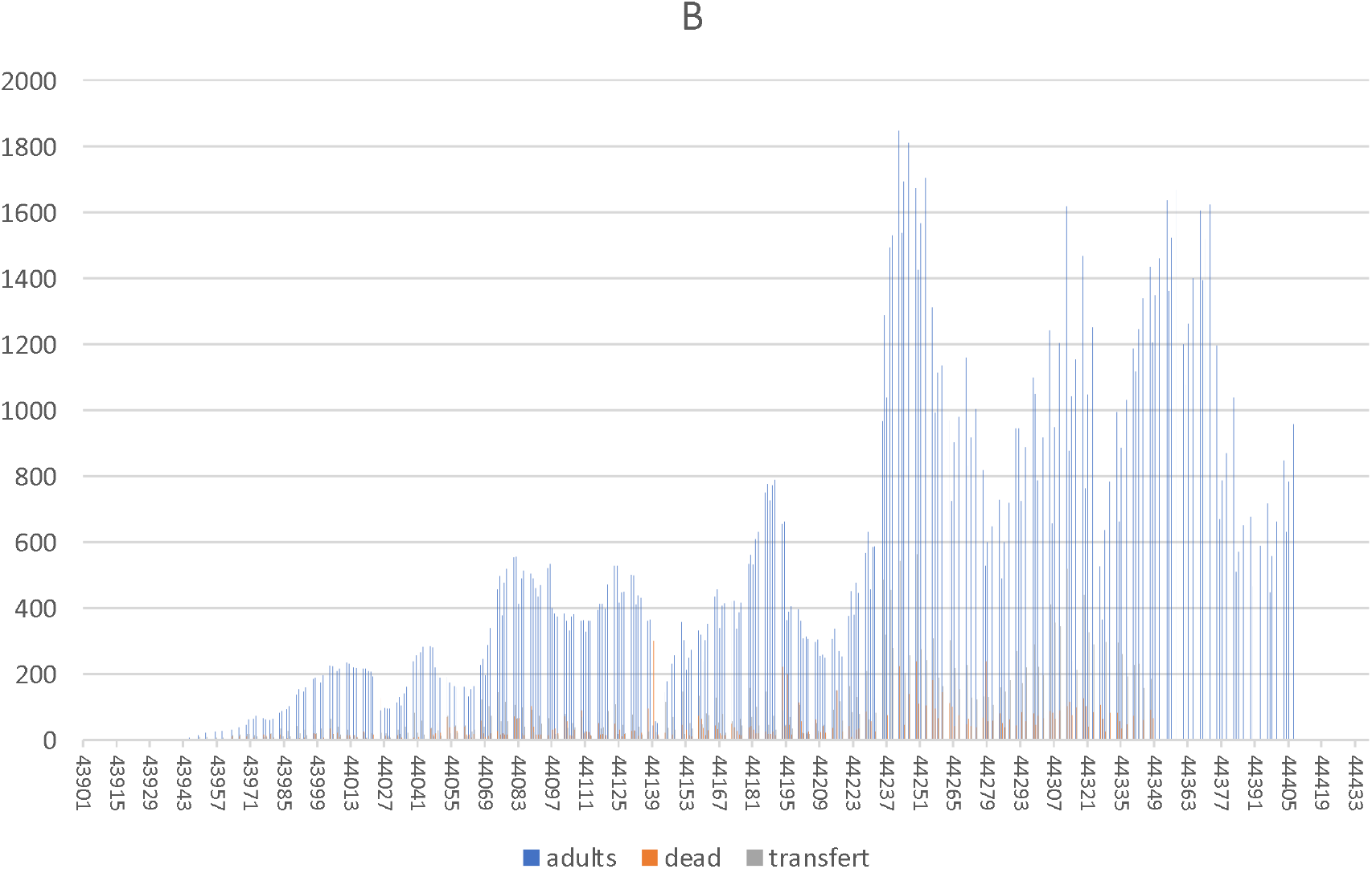
Number of individuals present, dead and transferred to the colony per day for *Phlebotomus perniciosus* (A) and *Phlebotomus papatasi* (B).

We were not able to take care of the sand flies every day during the COVID-19 lockdown period (March to May 2020), which led to a proliferation of fungi in the rearing pots and affected the mass production of sand flies. In addition, mites (Figure 2) invaded our colonies. These insects can feed on eggs, which also influences mass production.

**Figure 2.**
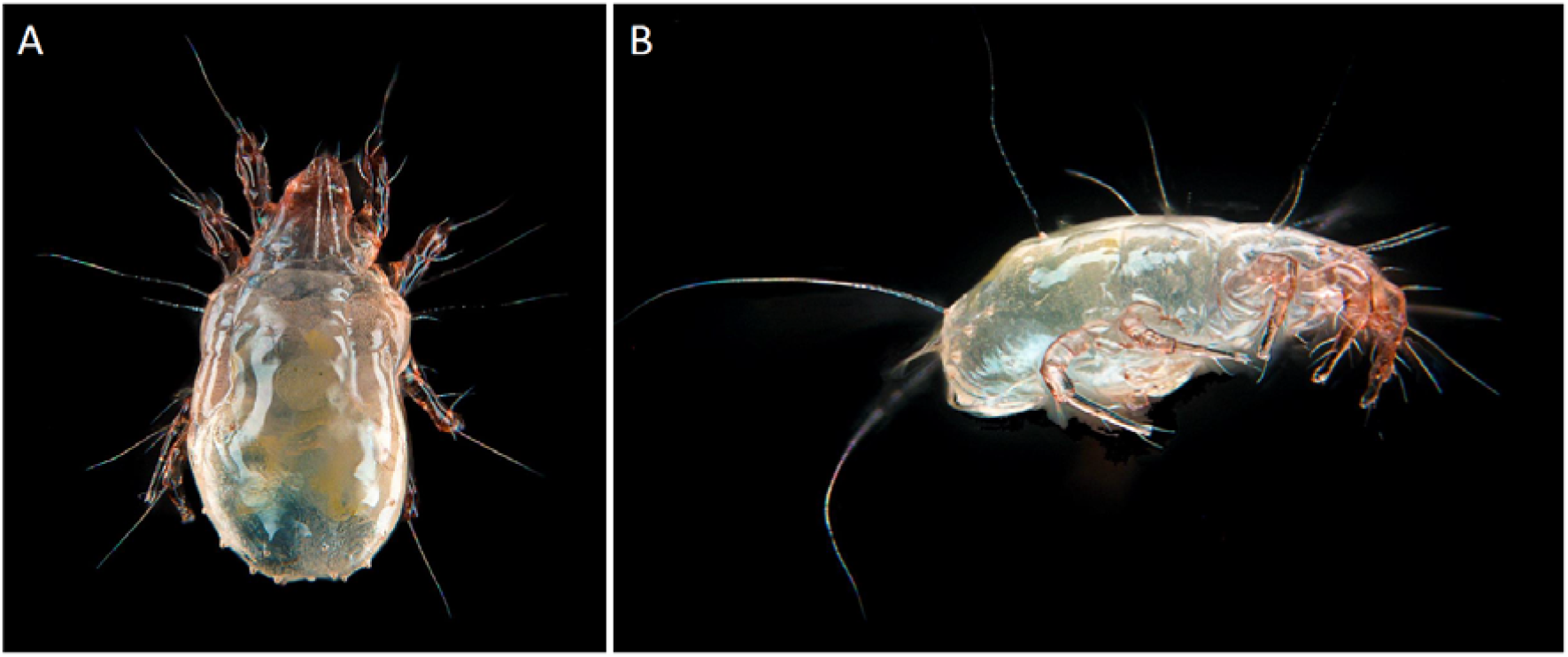
Phorid mites found in the colony. A. Dorsal aspect; B. Lateral aspect. (Photos by N. Rahola).

At the end of COVID-19 confinement measures, we managed to overcome the fungus problem. On the other hand, mite infestations are almost impossible to avoid, we are currently testing several processes to eliminate them (*e*.*g*., daily removal of dead individuals, ovipots checked daily, mite removal) (Lawyer P et al., 2017).

The establishment of the colonies and the survival rate over two years and several generations show that the membrane feeding system is viable and has multiple advantages:

- it avoids the use of live animals;
- it does not require trained personnel;
- it allows the choice of blood;
- it avoids colony adaptation to alternative blood, which can be a long process and can take several generations (Volf & Volfova, 2011).

### Colony productivity

Table 1 shows the productivity by generation. An average of all generations can be estimated. In total, 5,284 *Ph. papatasi* females and 10,651 *Ph. perniciosus* females were put in rearing pots. To date, the number of emerged adults was 24,773 and 38,435 and the mean number of females was 9.9 and 6.9 for *Ph. papatasi* and *Ph. perniciosus*, respectively.

We observed a decrease of productivity after the first generation for *Ph. perniciosus*. Among the many factors that can be responsible for this decrease, two explanations seem more likely. First, fungi were present in the rearing pots in the second generation, affecting larvae mobility and feeding behavior. Second, the colony may have needed to adapt to our insectary (*e*.*g*., climatic condition, manipulation). However, we observed an increase in productivity of *Ph. perniciosus* females at the fourth generation, which could be the result of a good adaptation to the insectary. However, there was a decrease at the seventh generation probably due to the female number per pot. Indeed, a high density of females can generate stress and lead to a decrease of eggs production (Lawyer P et al., 2017).

With a 50/50 sex ratio, the mean percentage of blood-fed females during blood meals is higher for *Ph. papatasi* (23% [1.93; 58.61]) than for *Ph. perniciosus* (21% [1.59; 61.36]).

For *Ph. papatasi* and *Ph. perniciosus*, 29% and 17% of females died before laying eggs, respectively. However, these results are probably underestimated because it was not possible to determine if all the females laid eggs after the first egg appearance. These data also highlighted that our *Ph. perniciosus* colony adapted well whereas our *Ph. papatasi* colony needed time to adapt.

### Colony weekly production

For *Ph. perniciosus*, we were able to conduct six experimental infections with 1951 females, with the first one performed 10 months after the establishment of the colony. We reached a stable weekly production of at least 800 individuals one year after the start of the colony. For *Ph. papatasi*, we used almost 6,000 individuals for insecticide resistance tests, with a first experimentation performed 11 months after the establishment of the colony. A stable weekly production of at least 400 individuals was reached 11 months after the start of the colony (Figure 3).

**Figure 3.**
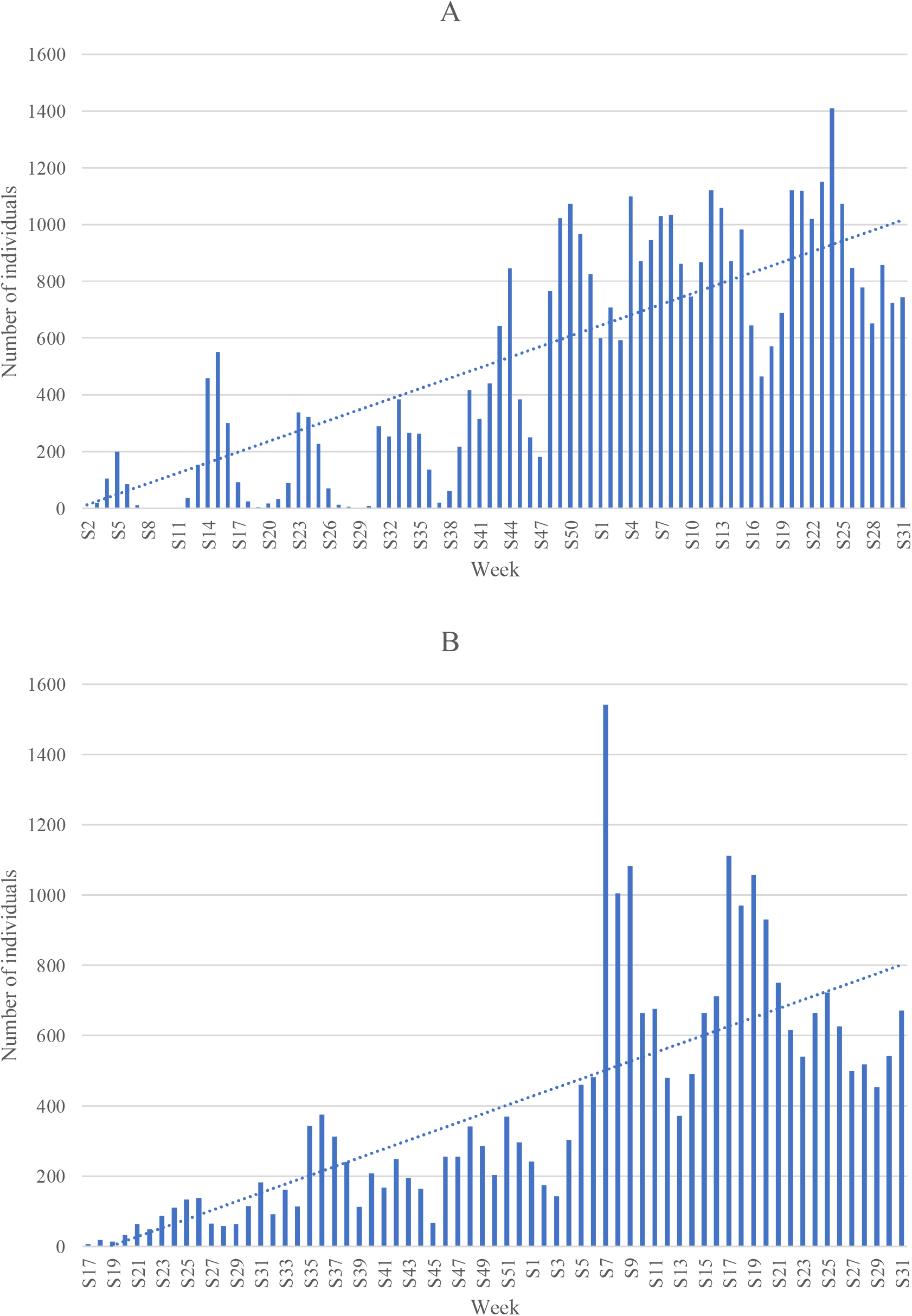
Number of individuals produced by week for *Phlebotomus perniciosus* (A) and *Phlebotomus papatasi* (B).

### Generation time

Mean periods for preoviposition, hatching, development of larvae and pupae, and for the total developmental time per generation are reported in Table 2. The total developmental time, from blood meal to the next generation of adult emergence, is 49.9 days [48.4; 51.1] and 47.4 days [43.7; 52.0], for *Ph. papatasi* and *Ph. perniciosus*, respectively. The data presented in this work are similar to the data found in the literature (Maroli et al., 1987; Campbell et al., 1989; Volf & Volfova, 2011; Lawyer P et al., 2017).

Hatching period and developmental times of larvae and pupae were not significantly different among the generations for the two species (*p-*value = 0.9808 and 0.9999 for *Ph. papatasi* and *Ph. perniciosus*, respectively).

As expected (Lawyer P et al., 2017), we observed semi-asynchronous emergence. The adults emerged over an average period of 38.5 [18; 34.9] and 61.9 [41.3; 73.1] days calculated on all generations for *Ph. perniciosus* and *Ph. papatasi*, respectively.

To date, we observe a lower number of larval diapauses for *Ph. perniciosus* than for *Ph. papatasi*, which could be the result of the better adaptation of *Ph. perniciosus* to our insectary (Campbell et al., 1989).

## Appendices

Detailed protocol and file are available in Appendix A: Detailed protocol for 50% glucose solution preparation, Appendix B: Detailed protocol for sterile dissection of chicken skins., Appendix C: Detailed protocol for larval food preparation, and Appendix D: Detailed file on standardized data collection and tracking of sand fly colonies.

## Acknowledgments

The authors wish to thank for their financial support. We are also particularly grateful to Pr. Ricardo Molina and Pr. Petr Volf for providing the sand fly eggs to start our colonies. Idris Mhaidi is supported by the IHU Méditerannée Infection. We thank Heidi Lançon for editing and proofreading the manuscript.

## Supplementary information availability

Supplementary information are available online: https://doi.org/10.5281/zenodo.8163691.

